## Supplementary figures and images for "Assessment by Matrix-Assisted Laser Desorption – Time of Flight Mass Spectrometry of the Diversity of Endophytes and Rhizobacteria Cultured from the Maize Microbiome"

### Supplemental Figure S1

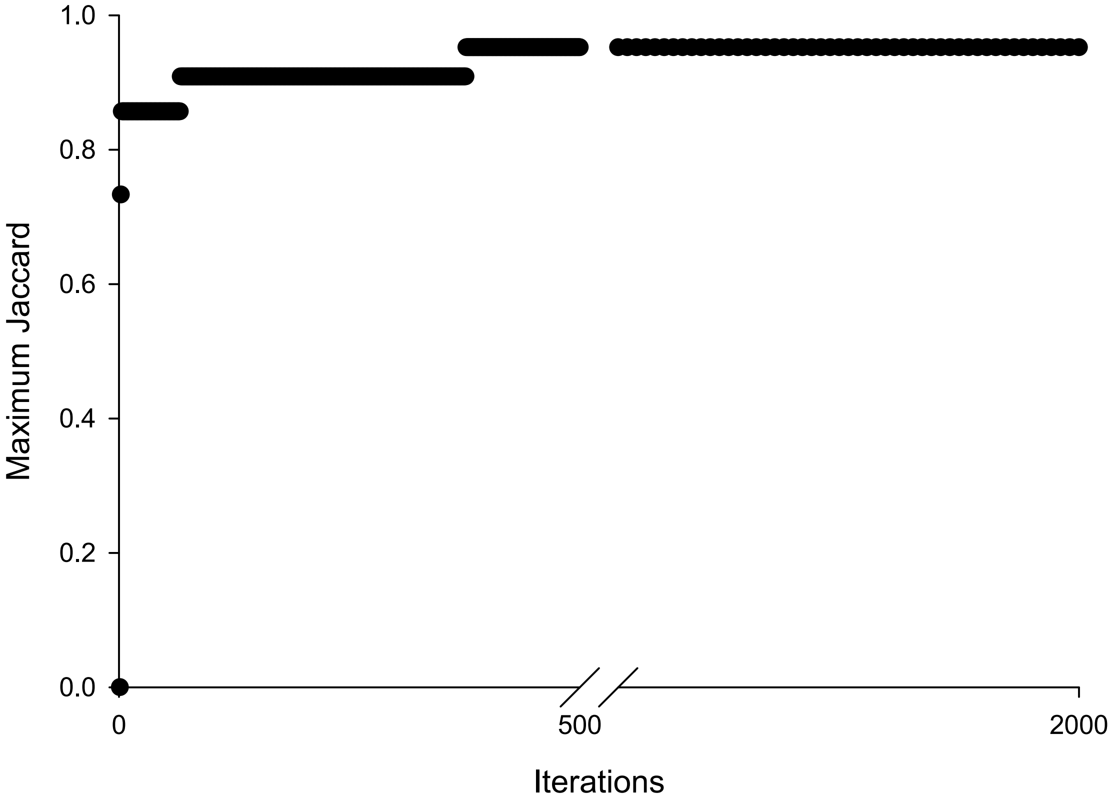

### Supplemental Figure S2

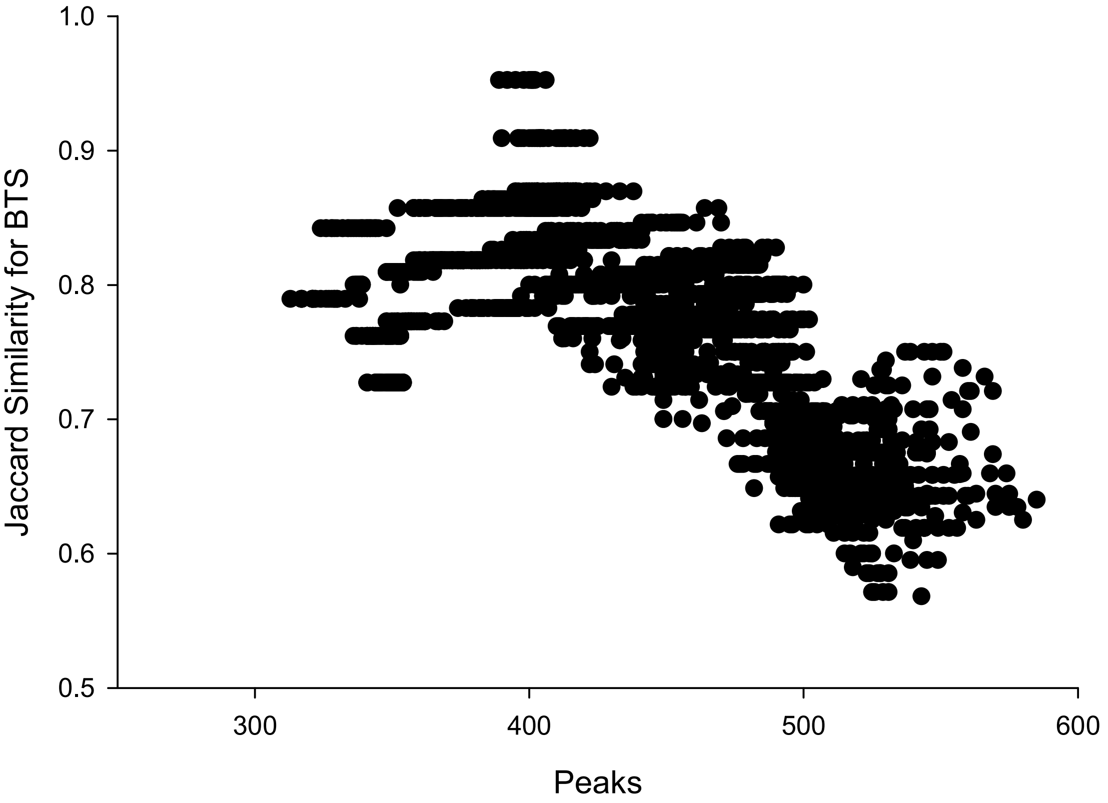

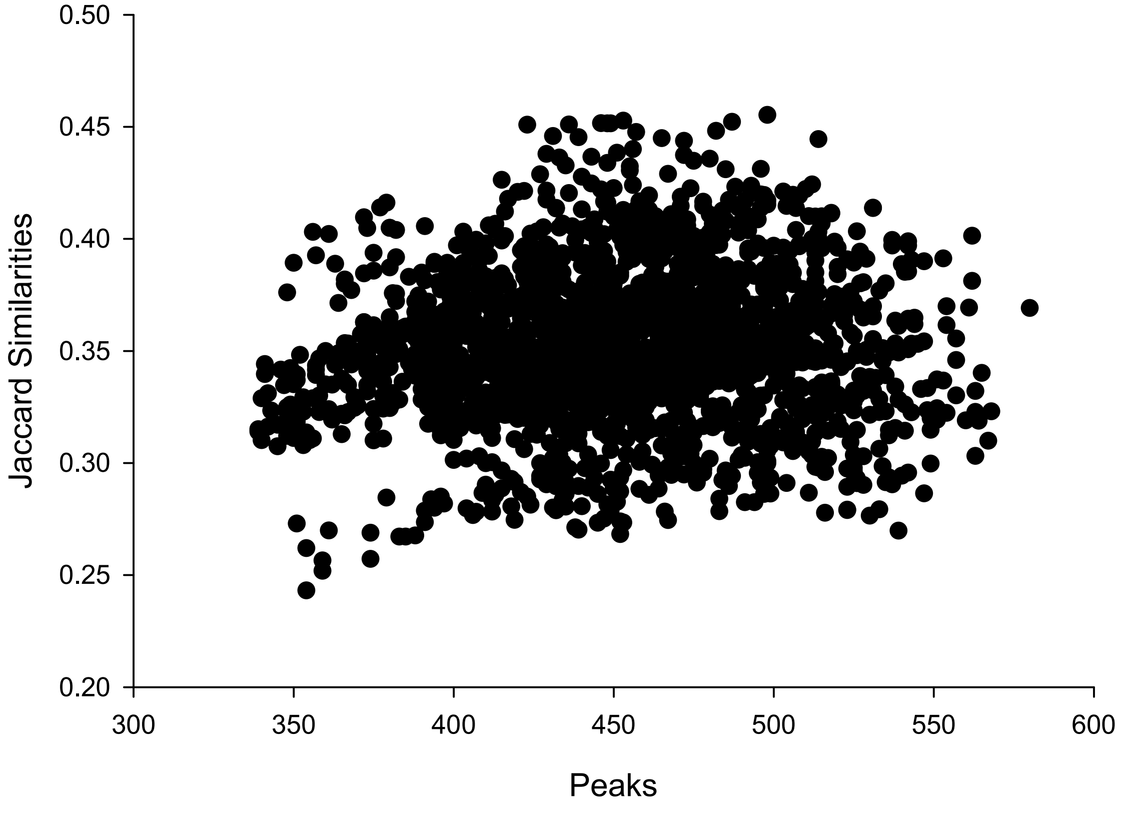

### Supplemental Figure S3

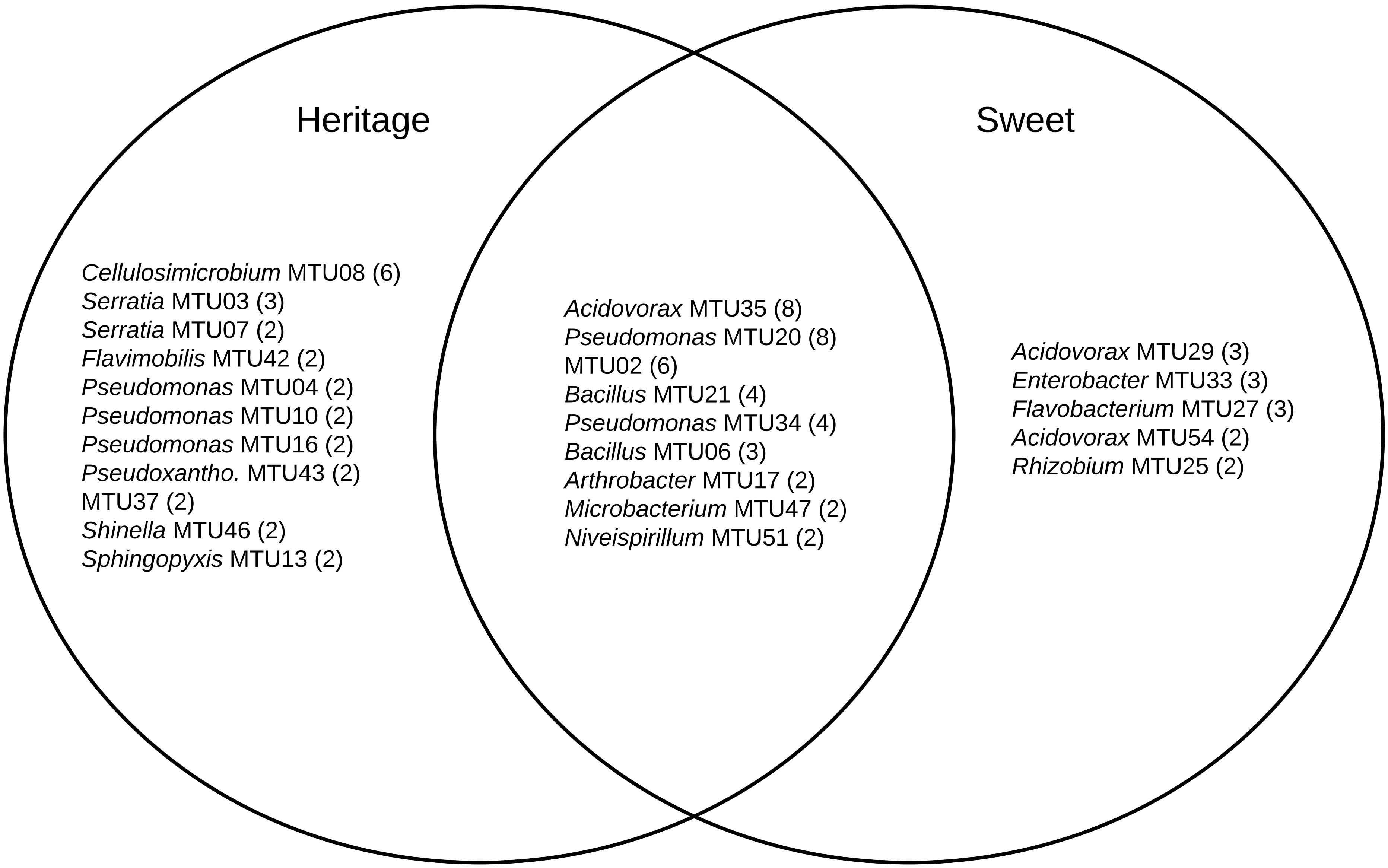
