## Supplementary material for "Assessment by Matrix-Assisted Laser Desorption – Time of Flight Mass Spectrometry of the Diversity of Endophytes and Rhizobacteria Cultured from the Maize Microbiome": Markdown File in html format: maize180308.html

Maize Microbiome


### Maize Microbiome

###### Michael G. LaMontagne

#### 11/27/2020

```
library("MALDIquant")
```

```
## 
## This is MALDIquant version 1.19.3
## Quantitative Analysis of Mass Spectrometry Data
##  See '?MALDIquant' for more information about this package.
```

```
library("MALDIquantForeign")
library("pvclust")
library(philentropy)
# library(ggplot2)
library(iNEXT)
library(RWeka)

packageVersion("MALDIquant")
```

```
## [1] '1.19.3'
```

```
packageVersion("MALDIquantForeign")
```

```
## [1] '0.12'
```

```
packageVersion("pvclust")
```

```
## [1] '2.2.0'
```

```
packageVersion("philentropy")
```

```
## [1] '0.4.0'
```

```
packageVersion("iNEXT")
```

```
## [1] '2.0.20'
```

```
packageVersion("ggplot2")
```

```
## [1] '3.3.2'
```

```
packageVersion("RWeka")
```

```
## [1] '0.4.42'
```

```
# write record of packages and versions
sessionInfo()
```

```
## R version 4.0.3 (2020-10-10)
## Platform: x86_64-w64-mingw32/x64 (64-bit)
## Running under: Windows 10 x64 (build 17763)
## 
## Matrix products: default
## 
## locale:
## [1] LC_COLLATE=English_United States.1252 
## [2] LC_CTYPE=English_United States.1252   
## [3] LC_MONETARY=English_United States.1252
## [4] LC_NUMERIC=C                          
## [5] LC_TIME=English_United States.1252    
## 
## attached base packages:
## [1] stats     graphics  grDevices utils     datasets  methods   base     
## 
## other attached packages:
## [1] RWeka_0.4-42           iNEXT_2.0.20           philentropy_0.4.0     
## [4] pvclust_2.2-0          MALDIquantForeign_0.12 MALDIquant_1.19.3     
## 
## loaded via a namespace (and not attached):
##  [1] Rcpp_1.0.4.6             readBrukerFlexData_1.8.5 plyr_1.8.6              
##  [4] compiler_4.0.3           pillar_1.4.6             RWekajars_3.9.3-2       
##  [7] base64enc_0.1-3          tools_4.0.3              digest_0.6.27           
## [10] evaluate_0.14            lifecycle_0.2.0          tibble_3.0.1            
## [13] gtable_0.3.0             pkgconfig_2.0.3          rlang_0.4.6             
## [16] yaml_2.2.1               parallel_4.0.3           xfun_0.19               
## [19] rJava_0.9-12             readMzXmlData_2.8.1      dplyr_0.8.5             
## [22] stringr_1.4.0            knitr_1.30               vctrs_0.3.0             
## [25] tidyselect_1.1.0         grid_4.0.3               glue_1.4.1              
## [28] R6_2.5.0                 XML_3.99-0.5             rmarkdown_2.5           
## [31] reshape2_1.4.4           purrr_0.3.4              ggplot2_3.3.2           
## [34] magrittr_1.5             scales_1.1.1             htmltools_0.5.0         
## [37] ellipsis_0.3.1           assertthat_0.2.1         colorspace_1.4-1        
## [40] stringi_1.4.6            munsell_0.5.0            crayon_1.3.4
```

#### If starting from raw spectra, import spectra from two runs. Files should be in folders with paths like this: C:-library\4.0\180308old\0\_A1\1\1SLin

```
# import test spectra 1 (not run in markdown)
#exampleDirectory <- #system.file("180308old",package="MALDIquantForeign")
#spectraRold <- import(file.path(exampleDirectory,"brukerflex#"),verbose=FALSE)
# import spectra 2
#exampleDirectory <- #system.file("180308new",package="MALDIquantForeign")
#spectraRnew <- import(file.path(exampleDirectory,"brukerflex#"),verbose=FALSE)
```

#### trim and transform raw spectra (skip in markdown)

```
# set range to trim outside of 
#range = c(1900, 12000)
# trim beginging and ending
#spectraOld2 <- MALDIquant::trim(spectraRold, range=range)
#spectraNew2 <- MALDIquant::trim(spectraRnew, range=range)
# transform
#spectraOld2 <- transformIntensity(spectraOld2, method="sqrt")
#spectraNew2 <- transformIntensity(spectraNew2, method="sqrt")
# check spectra
#head(spectraOld2)
#head(spectraNew2)
```

#### Merge two spectra files (if starting from raw spectra)

```
## merge and print total length
# spectraB <- append(spectraNew2, spectraOld2) 
# length(spectraB)
```

#### create reference peaks

```
# 
rR <- createMassPeaks(mass=c(2833, 4362, 5095, 5379, 5392, 5396, 6254,
                             6268, 6278, 6291, 6314, 6328, 6353, 6410, 7272, 9533, 9552),
                      intensity=rep(1, 17))
rR
```

```
## S4 class type            : MassPeaks    
## Number of m/z values     : 17           
## Range of m/z values      : 2833 - 9552  
## Range of intensity values: 1e+00 - 1e+00
## Range of snr values      : NA - NA      
## Memory usage             : 1.531 KiB
```

#### run optimize alignment loop (start here with example data)

```
spectraB <- readRDS("spectra180308b")
# create variables
y <- c(0, 0, 0, 0, 0, 0, 0, 0, 0, 0, 0, 0, 0)
Draw <- matrix(y, nrow=13, ncol=1)
rownames(Draw) <- c("hwSmth", "rmvbI", "hwAlgn", "snrAlgn", "tolAlgn", "hwDtct", "snrDtct", "peaks", "Cor12", "RichE", "K12", "Pass", "MaxRep")
# reset counts
try <- c(0)
Pass <- c(0)
M12 <- c(0)
# align test spectra against reference spectra 
repeat{
  hwSmth <- 10*(sample(7:10, 1))
  rmvbI <- 25*(sample(3:4, 1))
  hwA <-  10*(sample(5:9, 1))
  snrA <-  (sample(26:30, 1))/10
  tolA <- (sample(23:30, 1))/1000
  hwD <- 10*(sample(5:10, 1))
  snrD <- (sample(40:90, 1))/10
  ##align
  spectra <- smoothIntensity(spectraB, method="SavitzkyGolay", halfWindowSize=hwSmth)
  spectra <- removeBaseline(spectra, method="SNIP",iterations=rmvbI)
  spectraT <- calibrateIntensity(spectra, method="TIC")
  spectra <- try(alignSpectra 
                 (spectraT, halfWindowSize=hwA, SNR=snrA, reference = rR, 
                   tolerance=tolA, warpingMethod="lowess"), silent = FALSE)
if ('try-error' %in% class(spectra)) next
  ##---metadata
  LB  <- length(spectra)
  metaData(spectra[[LB]])$sampleName
  spots <- sapply(spectra, function(x)metaData(x)$spot)
  species <- sapply(spectra, function(x)metaData(x)$sampleName)
  avgSpectra <-
    averageMassSpectra((spectra), labels=paste0(species))
## detectPeaks
  try(peaks <- detectPeaks(avgSpectra, SNR=snrD, halfWindowSize=hwD), silent = FALSE)
  if ('try-error' %in% class(avgSpectra)) next
  ##--export peaks
  peaks <- binPeaks(peaks)
  spots <- sapply(avgSpectra, function(x)metaData(x)$spot)
  species <- sapply(avgSpectra, function(x)metaData(x)$sampleName)
  species <- factor(species)
  n <- length(avgSpectra) 
  featureMatrix <- intensityMatrix(peaks, avgSpectra)
  rownames(featureMatrix) <- paste(species)
  peaksNew <- intensityMatrix(peaks)
  rownames(peaksNew) <- paste(species)
  ##--write matrix
  FMtrnsp <- t(featureMatrix)
  nrow(FMtrnsp)
  # calculate similarity matrix
  x <- cor(FMtrnsp)
  rownames(x) <- paste(species)
  # calcualte cos simmilarity
  simCos <- coop::cosine(FMtrnsp)
  rownames(simCos) <- paste(species)
  ## extract correlations within BTS and 
  Cor12 <- (simCos[1,65])
  ## count pass
  Pass <- Pass + 1
  ## calculate Jaccard coefficients
  N <- intensityMatrix(peaks)
  rownames(N) <- paste(species)
  N[N>0] <- 1
  N[is.na(N)] <- 0
  k <- distance((N), method = "jaccard")
  Rich <- ncol(N)
  richE <- sum(N[1,])
  # calculate BTS match
  K12 <- 1-(k[1,65])
  ## name each data value
  SNRalgn <- c(hwSmth, rmvbI, hwA, snrA, tolA, hwD, snrD, Rich, Cor12, richE, K12, Pass, M12)
  names(SNRalgn) <- c("hwSmth", "rmvbI", "hwAlgn", "snrAlgn", "tolAlgn", "hwDtct", "snrDtct", "peaks", "Cor12", "RichE", "K12", "Pass", "MaxRep")
  B = matrix(SNRalgn, nrow=13, ncol=1) 
  Draw <- cbind(B, Draw)
  M12 <- max(Draw[11,])
# set tries to 40 for markdown, 2000 tries used to generate 
  # D180308raw
  if (Pass > 40) {break}
}# end of align loop
```

```
## Error in FUN(X[[i]], ...) : 
##   Could not match any peak in spectrum 55 to a reference peak.
## Error in FUN(X[[i]], ...) : 
##   Could not match enough peaks in spectrum62to the reference peaks.
## 2 matches required, just 1 found.
## Error in FUN(X[[i]], ...) : 
##   Could not match enough peaks in spectrum62to the reference peaks.
## 2 matches required, just 1 found.
## Error in FUN(X[[i]], ...) : 
##   Could not match enough peaks in spectrum55to the reference peaks.
## 2 matches required, just 1 found.
## Error in FUN(X[[i]], ...) : 
##   Could not match enough peaks in spectrum55to the reference peaks.
## 2 matches required, just 1 found.
## Error in FUN(X[[i]], ...) : 
##   Could not match enough peaks in spectrum55to the reference peaks.
## 2 matches required, just 1 found.
## Error in FUN(X[[i]], ...) : 
##   Could not match enough peaks in spectrum62to the reference peaks.
## 2 matches required, just 1 found.
## Error in FUN(X[[i]], ...) : 
##   Could not match enough peaks in spectrum55to the reference peaks.
## 2 matches required, just 1 found.
## Error in FUN(X[[i]], ...) : 
##   Could not match enough peaks in spectrum62to the reference peaks.
## 2 matches required, just 1 found.
## Error in FUN(X[[i]], ...) : 
##   Could not match enough peaks in spectrum55to the reference peaks.
## 2 matches required, just 1 found.
## Error in FUN(X[[i]], ...) : 
##   Could not match enough peaks in spectrum55to the reference peaks.
## 2 matches required, just 1 found.
## Error in FUN(X[[i]], ...) : 
##   Could not match enough peaks in spectrum62to the reference peaks.
## 2 matches required, just 1 found.
## Error in FUN(X[[i]], ...) : 
##   Could not match enough peaks in spectrum55to the reference peaks.
## 2 matches required, just 1 found.
## Error in FUN(X[[i]], ...) : 
##   Could not match enough peaks in spectrum55to the reference peaks.
## 2 matches required, just 1 found.
## Error in FUN(X[[i]], ...) : 
##   Could not match enough peaks in spectrum101to the reference peaks.
## 2 matches required, just 1 found.
## Error in FUN(X[[i]], ...) : 
##   Could not match any peak in spectrum 55 to a reference peak.
## Error in FUN(X[[i]], ...) : 
##   Could not match enough peaks in spectrum55to the reference peaks.
## 2 matches required, just 1 found.
## Error in FUN(X[[i]], ...) : 
##   Could not match enough peaks in spectrum62to the reference peaks.
## 2 matches required, just 1 found.
## Error in FUN(X[[i]], ...) : 
##   Could not match enough peaks in spectrum62to the reference peaks.
## 2 matches required, just 1 found.
## Error in FUN(X[[i]], ...) : 
##   Could not match enough peaks in spectrum55to the reference peaks.
## 2 matches required, just 1 found.
## Error in FUN(X[[i]], ...) : 
##   Could not match enough peaks in spectrum62to the reference peaks.
## 2 matches required, just 1 found.
## Error in FUN(X[[i]], ...) : 
##   Could not match enough peaks in spectrum55to the reference peaks.
## 2 matches required, just 1 found.
## Error in FUN(X[[i]], ...) : 
##   Could not match enough peaks in spectrum55to the reference peaks.
## 2 matches required, just 1 found.
```

```
# saveRDS(D, "D180308raw")
# write.csv(as.matrix(t(Draw)), "Draw.csv")
```

#### plot jaccard similarity versus try

```
# import Draw generated with 2000 iterations
Draw <- readRDS("D180308raw")
plot.default(Draw[12,], Draw[13,], xlab = "Try", ylab = "Jaccard")
```

#### plot jaccard similarity versus peaks

```
plot.default(Draw[8,], Draw[11,], xlab = "Peaks", ylab = "Jaccard")
```

```
# write.csv(k, "kRef.csv", row.names = rownames(N))
# write.csv(simCos, "simCosRef.csv")
```

#### select optimized parameters

```
DF <- as.data.frame(t(Draw))
DF <- DF[order(-DF$K12, -DF$peaks),]
dim(DF)
```

```
## [1] 2002   13
```

```
hwSmth <- DF$hwSmth[1]
rmvbI <- DF$rmvbI[1]
hwA <-  DF$hwA[1]
snrA <-  DF$snrA[1]
tolA <- DF$tolA[1]
hwD <- DF$hwD[1]
snrD <- DF$snrD[1]
hwSmth
```

```
## [1] 80
```

```
rmvbI
```

```
## [1] 75
```

```
hwA
```

```
## [1] 80
```

```
snrA
```

```
## [1] 2.9
```

```
tolA
```

```
## [1] 0.03
```

```
hwD
```

```
## [1] 50
```

```
snrD
```

```
## [1] 9
```

#### generate feature matrix with optimized

```
##align spectra
spectra <- smoothIntensity(spectraB, method="SavitzkyGolay", halfWindowSize=hwSmth)
spectra <- removeBaseline(spectra, method="SNIP",iterations=rmvbI)
spectraT <- calibrateIntensity(spectra, method="TIC")
spectra <- (alignSpectra (spectraT, halfWindowSize=hwA, SNR=snrA, reference = rR, tolerance=tolA, warpingMethod="lowess"))
##---metadata
spots <- sapply(spectra, function(x)metaData(x)$spot)
list(spots)
```

```
## [[1]]
##   [1] "A1"  "A2"  "A5"  "A8"  "A6"  "A9"  "A10" "A7"  "A11" "A12" "B1"  "B3" 
##  [13] "B5"  "B6"  "B7"  "C7"  "C8"  "C10" "C9"  "F7"  "F8"  "F9"  "F10" "H10"
##  [25] "H9"  "H11" "C11" "C12" "D1"  "D2"  "C5"  "D3"  "D4"  "D5"  "D6"  "D7" 
##  [37] "D8"  "D9"  "F12" "G1"  "G2"  "G3"  "B8"  "B9"  "B10" "B11" "B12" "D10"
##  [49] "D11" "D12" "E2"  "E3"  "E4"  "E5"  "E9"  "E10" "E6"  "E11" "E7"  "E12"
##  [61] "E8"  "F1"  "F2"  "F3"  "F4"  "G7"  "G8"  "G10" "G11" "G12" "C2"  "C3" 
##  [73] "C4"  "F5"  "H1"  "H2"  "H3"  "H4"  "H6"  "H7"  "A1"  "A2"  "H11" "H12"
##  [85] "H3"  "H4"  "H5"  "H6"  "H7"  "H8"  "H10" "H9"  "A5"  "A6"  "A7"  "A8" 
##  [97] "A9"  "A10" "A11" "A12" "B1"  "B2"  "B3"  "B4"  "B5"  "B6"  "G5"  "G6" 
## [109] "G7"  "G8"  "G10" "G9"  "B8"  "B9"  "B10" "B11" "B12" "C1"  "C2"  "C3" 
## [121] "C4"  "C5"  "C6"  "B7"  "C7"  "C8"  "C10" "C9"  "C11" "C12" "D1"  "D2" 
## [133] "F11" "F12" "G1"  "G2"  "G3"  "G4"  "G11" "G12" "H1"  "H2"  "D3"  "D4" 
## [145] "D5"  "D6"  "D7"  "D8"  "D10" "D9"  "D11" "D12" "E1"  "E2"  "E3"  "E4" 
## [157] "E5"  "E6"  "E7"  "E8"  "E10" "E9"  "E11" "E12" "F1"  "F2"  "F3"  "F4" 
## [169] "F5"  "F6"  "F7"  "F8"  "F10" "F9"
```

```
species <- sapply(spectra, function(x)metaData(x)$sampleName)
list(species)
```

```
## [[1]]
##   [1] "BTSnew"  "BTSnew"  "H1Ne01"  "H1Ne01"  "H1Ne02"  "H1Ne02"  "H1Ne03" 
##   [8] "H1Ne03"  "H1Ne04"  "H1Ne05"  "H1Ne06"  "H1Ne08"  "H1Ne10"  "H1Ne11" 
##  [15] "H1Ne12"  "H1Nz23"  "H1Nz25"  "H1Nz26"  "H1Nz26"  "H1Oe55"  "H1Oe56" 
##  [22] "H1Oe57"  "H1Oe58"  "H1Oz81"  "H1Oz81"  "H1Oz83"  "H2Nz28"  "H2Nz29" 
##  [29] "H2Nz30"  "H2Nz30"  "H3Ne22"  "H3Nz32"  "H3Nz33"  "H3Nz34"  "H3Nz35" 
##  [36] "H3Nz36"  "H3Nz37"  "H3Nz38"  "H3Oe60"  "H3Oe61"  "H3Oe62"  "H3Oe63" 
##  [43] "S1Ne13"  "S1Ne14"  "S1Ne15"  "S1Ne16"  "S1Ne17"  "S1Nz39"  "S1Nz40" 
##  [50] "S1Nz40"  "S1Nz42"  "S1Nz42"  "S1Nz44"  "S1Nz45"  "S1Nz45"  "S1Nz46" 
##  [57] "S1Nz46"  "S1Nz47"  "S1Nz47"  "S1Nz48"  "S1Nz48"  "S1Nz49"  "S1Nz50" 
##  [64] "S1Nz51"  "S1Nz52"  "S1Oe67"  "S1Oe67"  "S1Oe70"  "S1Oe70"  "S1Oe72" 
##  [71] "S3Ne19"  "S3Ne19"  "S3Ne21"  "S3Nz53"  "S4Oe73"  "S4Oe74"  "S4Oe75" 
##  [78] "S4Oe76"  "S4Oe78"  "S4Oe79"  "BTSold"  "BTSold"  "BTSold"  "BTSold" 
##  [85] "H1NZ141" "H1NZ141" "H1NZ142" "H1NZ142" "H1NZ143" "H1NZ143" "H1NZ144"
##  [92] "H1NZ144" "H1OZ85"  "H1OZ86"  "H1OZ87"  "H1OZ88"  "H1OZ89"  "H1OZ90" 
##  [99] "H2OZ91"  "H2OZ92"  "H2OZ93"  "H2OZ94"  "H2OZ95"  "H2OZ96"  "H2OZ97" 
## [106] "H2OZ98"  "H3NZ135" "H3NZ135" "H3NZ137" "H3NZ137" "H3NZ138" "H3NZ138"
## [113] "H3OZ100" "H3OZ101" "H3OZ102" "H3OZ103" "H3OZ104" "H3OZ105" "H3OZ105"
## [120] "H3OZ106" "H3OZ106" "H3OZ107" "H3OZ107" "H3OZ99"  "H4OZ109" "H4OZ109"
## [127] "H4OZ110" "H4OZ110" "H4OZ111" "H4OZ111" "H4OZ112" "H4OZ112" "S1NZ132"
## [134] "S1NZ132" "S1NZ133" "S1NZ133" "S1NZ134" "S1NZ134" "S2NZ139" "S2NZ139"
## [141] "S2NZ140" "S2NZ140" "S2OZ113" "S2OZ113" "S2OZ114" "S2OZ114" "S2OZ115"
## [148] "S2OZ115" "S2OZ116" "S2OZ116" "S2OZ117" "S2OZ117" "S2OZ120" "S2OZ120"
## [155] "S3OZ121" "S3OZ121" "S3OZ122" "S3OZ122" "S3OZ123" "S3OZ123" "S4OZ124"
## [162] "S4OZ124" "S4OZ125" "S4OZ125" "S4OZ126" "S4OZ126" "S4OZ127" "S4OZ127"
## [169] "S4OZ128" "S4OZ128" "S4OZ129" "S4OZ129" "S4OZ130" "S4OZ130"
```

```
species <- factor(species)
## detectPeaks
peaks <- detectPeaks(spectra, SNR=snrD, halfWindowSize=hwD)
# reset matrix
x <- c(0.1:0.6)
E <- matrix(x, nrow=6, ncol=1)
rownames(E) <- c("speciesE", "tryE", "noiseE", "signal", "SNR", "peaks")
##
n <- length(spectra)
n
```

```
## [1] 174
```

```
# saveRDS(spectra, "spectra180308raw")
```

#### run SNR loop

```
# to restart here
# spectra <- readRDS("spectra180308raw")
tryE <-c(0)
repeat{
  tryE <- tryE + 1 
# view one particular spectra 
# tryE <- 63
  noise <- estimateNoise(spectra[[tryE]])
#  plot(spectra[[tryE]], xlim=c(2000, 12000), ylim=c(0, 0.002))
#  points(peaks[[tryE]], col="red", pch=4)
  
##--generate matrix
featureMatrix <- intensityMatrix(peaks, spectra)
  rownames(featureMatrix) <- paste(species, spots)
  nrow(featureMatrix)
  ## calculate Signal
  noise <- estimateNoise(spectra[[tryE]])
  head(noise)
speciesE <- rownames(featureMatrix)
speciesE <- speciesE[tryE]
  EaPeaks <- intensityMatrix(peaks[tryE])
  richE <- ncol(EaPeaks)
  sorted <- EaPeaks[order(EaPeaks, decreasing = TRUE)]
  signal <- median((sorted[1:10]))
  noiseE <- noise[1, 2]
  SNRe <- (signal/noiseE)
  ## name each data value
  SNRall <- c(speciesE, tryE, noiseE, signal, SNRe, richE)
  names(SNRall) <- c("speciesE", "tryE", "noiseE", "signal", "SNR", "peaks")
  C = matrix(SNRall, nrow=6, ncol=1) 
  E <- cbind(C, E)
  if (tryE > (n-1)) {break}
}
plot(spectra[[tryE]], xlim=c(2000, 12000), ylim=c(0, 0.002))
points(peaks[[tryE]], col="red", pch=4)
```

#### plot SNR vs Peaks and export SNR

```
F <- t(E[,1:n])
F[is.na(F)] = 0
F <- as.data.frame(F)
F$tryE <- as.numeric(F$tryE)
F$SNR <- as.numeric(F$SNR)
F$peaks <- as.numeric(F$peaks)
# write.csv(as.matrix(F), "F.csv")
plot.default(F$SNR, F$peaks, xlab = "SNR", ylab = "peaks")
```

#### subset and count noisey spectra

```
Fnoisey <- F[ which(F$SNR < 15 & F$peaks < 13),]
# pull out specta numbers and sort them
noiseySpec <- c(Fnoisey$tryE)
noiseySpec <- sort(noiseySpec)
n <- length(noiseySpec)
n
```

```
## [1] 8
```

```
# saveRDS(noiseySpec, "noiseySpec180308")
# noiseySpec <- readRDS("noiseySpec180308")
```

#### plot noisey spectra

```
tryE <- c(0)
n <- length(noiseySpec)
n
```

```
## [1] 8
```

```
# run SNR loop
spectraN <- spectra[noiseySpec]
peaksN <- peaks[noiseySpec]
repeat{
  tryE <- tryE + 1 
  plot(spectraN[[tryE]], xlim=c(2000, 12000), ylim=c(0, 0.002))
  points(peaksN[[tryE]], col="red", pch=4)
  if (tryE > (n-1)) {break}}
```

#### subset quality spectra

```
# input file
# spectraB <- readRDS("spectra180308b")
spectraQ <- spectraB [-(noiseySpec)]
length(spectraQ)
```

```
## [1] 166
```

```
saveRDS(spectraQ, "spectra180308q")
# spectraQ <- readRDS("spectra180308q")
head(spectraQ)
```

```
## [[1]]
## S4 class type            : MassSpectrum         
## Number of m/z values     : 36211                
## Range of m/z values      : 1900.149 - 11999.606 
## Range of intensity values: 4.472e+00 - 7.934e+01
## Memory usage             : 576.18 KiB           
## Name                     : BTSnew.A1            
## File                     : C:\Users\lamontagne\Documents\R\win-library\4.0\MALDIquantForeign\180308new\brukerflex\BTSnew\0_A1\1\1SLin\fid
## 
## [[2]]
## S4 class type            : MassSpectrum         
## Number of m/z values     : 36211                
## Range of m/z values      : 1900.149 - 11999.606 
## Range of intensity values: 3.873e+00 - 8.543e+01
## Memory usage             : 576.18 KiB           
## Name                     : BTSnew.A2            
## File                     : C:\Users\lamontagne\Documents\R\win-library\4.0\MALDIquantForeign\180308new\brukerflex\BTSnew\0_A2\1\1SLin\fid
## 
## [[3]]
## S4 class type            : MassSpectrum         
## Number of m/z values     : 36211                
## Range of m/z values      : 1900.149 - 11999.606 
## Range of intensity values: 5.831e+00 - 1.546e+02
## Memory usage             : 576.18 KiB           
## Name                     : H1Ne01.A5            
## File                     : C:\Users\lamontagne\Documents\R\win-library\4.0\MALDIquantForeign\180308new\brukerflex\H1Ne01\0_A5\1\1SLin\fid
## 
## [[4]]
## S4 class type            : MassSpectrum         
## Number of m/z values     : 36211                
## Range of m/z values      : 1900.149 - 11999.606 
## Range of intensity values: 7.616e+00 - 1.977e+02
## Memory usage             : 576.18 KiB           
## Name                     : H1Ne01.A8            
## File                     : C:\Users\lamontagne\Documents\R\win-library\4.0\MALDIquantForeign\180308new\brukerflex\H1Ne01\0_A8\1\1SLin\fid
## 
## [[5]]
## S4 class type            : MassSpectrum         
## Number of m/z values     : 36211                
## Range of m/z values      : 1900.149 - 11999.606 
## Range of intensity values: 6.557e+00 - 1.322e+02
## Memory usage             : 576.18 KiB           
## Name                     : H1Ne02.A6            
## File                     : C:\Users\lamontagne\Documents\R\win-library\4.0\MALDIquantForeign\180308new\brukerflex\H1Ne02\0_A6\1\1SLin\fid
## 
## [[6]]
## S4 class type            : MassSpectrum         
## Number of m/z values     : 36211                
## Range of m/z values      : 1900.149 - 11999.606 
## Range of intensity values: 3.606e+00 - 1.165e+02
## Memory usage             : 576.18 KiB           
## Name                     : H1Ne02.A9            
## File                     : C:\Users\lamontagne\Documents\R\win-library\4.0\MALDIquantForeign\180308new\brukerflex\H1Ne02\0_A9\1\1SLin\fid
```

#### import 16S identity

```
# import 16S
RNAident <- read.table("16sIDmatrix.csv", sep=",", header = TRUE, check.names = TRUE, row.names = 1)
xy <- t(combn(colnames(RNAident), 2))
xyR <- data.frame(xy, dist=RNAident[xy])
xyR$rPair <- paste(xyR$X1,xyR$X2,sep="-")
names(xyR)[3] <- "distR"
# write.csv(as.matrix(xyR), "xyR.csv")
head(xyR)
```

```
##       X1      X2 distR          rPair
## 1 H1OZ89  H1OZ90 0.876  H1OZ89-H1OZ90
## 2 H1OZ89  H2OZ93 0.872  H1OZ89-H2OZ93
## 3 H1OZ89  H2OZ97 0.863  H1OZ89-H2OZ97
## 4 H1OZ89  H2OZ98 0.879  H1OZ89-H2OZ98
## 5 H1OZ89 H3NZ135 0.868 H1OZ89-H3NZ135
## 6 H1OZ89 H3NZ137 0.862 H1OZ89-H3NZ137
```

#### reset variables and import quality spectra

```
#input spectra if restarting here
# spectraQ <- readRDS("spectra180308q")
# create variables
y <- c(0, 0, 0, 0, 0, 0, 0, 0, 0, 0, 0, 0, 0, 0, 0, 0, 0, 0, 0, 0, 0, 0, 0)
Dq <- matrix(y, nrow=23, ncol=1)
rownames(Dq) <- c("hwSmth", "rmvbI", "hwAlgn", "snrAlgn", "tolAlgn", "hwDtct", "snrDtct", "peaks", "Cor12", "RichE", "K12", "Pass", "MaxRep", "fit16S", "out16S", "fitTry","fit16W", "out16W", "Km", "y0", "distJ", "distS", "distW")
# reset counts
try <- c(0)
Pass <- c(0)
M12 <- c(0)
```

#### align quality spectra

```
##align
# align test spectra against reference spectra 
repeat{
  hwSmth <- 10*(sample(5:9, 1))
  rmvbI <- 25*(sample(3:4, 1))
  hwA <-  10*(sample(4:9, 1))
  snrA <-  (sample(24:30, 1))/10
  tolA <- (sample(23:30, 1))/1000
  hwD <- 10*(sample(4:10, 1))
  snrD <- (sample(50:90, 1))/10
  ##align
  spectra <- smoothIntensity(spectraQ, method="SavitzkyGolay", halfWindowSize=hwSmth)
  spectra <- removeBaseline(spectra, method="SNIP",iterations=rmvbI)
  spectraT <- calibrateIntensity(spectra, method="TIC")
  spectra <- try(alignSpectra 
                 (spectraT, halfWindowSize=hwA, SNR=snrA, reference = rR, 
                   tolerance=tolA, warpingMethod="lowess"), silent = FALSE)
  if ('try-error' %in% class(spectra)) next
  ##---metadata
  LB  <- length(spectra)
  metaData(spectra[[LB]])$sampleName
  spots <- sapply(spectra, function(x)metaData(x)$spot)
  species <- sapply(spectra, function(x)metaData(x)$sampleName)
  avgSpectra <-
    averageMassSpectra((spectra), labels=paste0(species))
  ## detectPeaks
  try(peaks <- detectPeaks(avgSpectra, SNR=snrD, halfWindowSize=hwD), silent = FALSE)
  if ('try-error' %in% class(avgSpectra)) next
  ##--export peaks
  peaks <- binPeaks(peaks)
  spots <- sapply(avgSpectra, function(x)metaData(x)$spot)
  species <- sapply(avgSpectra, function(x)metaData(x)$sampleName)
  species <- factor(species)
  n <- length(avgSpectra) 
  featureMatrix <- intensityMatrix(peaks, avgSpectra)
  rownames(featureMatrix) <- paste(species)
  peaksNew <- intensityMatrix(peaks)
  rownames(peaksNew) <- paste(species)
  ##--write matrix
  FMtrnsp <- t(featureMatrix)
  nrow(FMtrnsp)
  # calculate similarity matrix
  x <- cor(FMtrnsp)
  rownames(x) <- paste(species)
  # calcualte cos simmilarity
  simCos <- coop::cosine(FMtrnsp)
  rownames(simCos) <- paste(species)
  ## extract correlations within BTS and 
  Cor12 <- (simCos[1,61])
  ## count pass
  Pass <- Pass + 1
  ## calculate Jaccard coefficients
  N <- intensityMatrix(peaks)
  rownames(N) <- paste(species)
  N[N>0] <- 1
  N[is.na(N)] <- 0
  k <- distance((N), method = "jaccard")
  Rich <- ncol(N)
  richE <- sum(N[1,])
  # calculate BTS match
  K12 <- 1-(k[1,61])
# create Jaccard similarity
rownames(k) <- paste(species)
colnames(k) <- paste(species)
k2 <- 1 - k
# create dataframe of simCos pairwise comparisons
  xySC <- t(combn(colnames(simCos), 2))
  xyS <- data.frame(xySC, dist=simCos[xySC])
  names(xyS)[3] <- "distS"
  xyS$sPair <- paste(xyS$X1,xyS$X2,sep="-")
#  head(xyS)
  xyJC <- t(combn(colnames(k2), 2))
  xyJ <- data.frame(xyJC, distJ=k2[xyJC])
#  head(xyJ)
  ## merge jaccard and simCos
  xyJS <- data.frame(xyS,xyJ[3])
# calculate Yo
y0 <- xyJS[ which(xyJS$distJ < 0.01),]
# hist(y0$distS)
y0 <- exp(mean(log(y0$distS)))
Km <- 0.2
# plot predicted simCos
xyJS$cosPred <- y0 + (xyJS$distJ/(Km+xyJS$distJ))
# plot.default(xyJS$distJ, xyJS$cosPred, xlab = "Jaccard", ylab = "SimCos")
# plot.default(xyJS$distS, xyJS$cosPred, xlab = "Jaccard", ylab = "SimCos")
# predict matrix
kW <- y0 + (k2/(Km+k2))
# weigh matrix
cosW <- (simCos + kW)/2

# create dataframe of weighted pairwise comparisons
  xyW <- t(combn(colnames(cosW), 2))
  xyW <- data.frame(xyW, distCW=cosW[xySC])
  names(xyW)[3] <- "distW"
  xyS <- data.frame(xyJS, xyW[3]) 
  head(xyS)
# write.csv(as.matrix(xyS), "xyS.csv")
# merge 16S and simCos
mRM <- merge(xyR, xyS, by, by.x="rPair", by.y="sPair", sort = TRUE)
# write.csv(as.matrix(mRM), "mRMw.csv")
# calculate mass spec similarity within species
mRM16S <- mRM[ which(mRM$distR > 0.99),]
# hist(mRM16S$distS)
# hist(mRM16S$distW)
fit16S <- min(mRM16S$distS)
fit16W <- min(mRM16S$distW)
distJ <- mean(mRM16S$distJ)
distS <- mean(mRM16S$distS)
distW <- mean(mRM16S$distW)
# calculate mass spec similarity between phyla
mRMp <- mRM[ which(mRM$distR < 0.9),]
# hist(mRMp$distS)
out16S <- max(mRMp$distS) 
out16W <- max(mRMp$distW)
fitTry <- fit16W - out16W
  ## name each data value
  SNRalgn <- c(hwSmth, rmvbI, hwA, snrA, tolA, hwD, snrD, Rich, Cor12, richE, K12, Pass, M12, fit16S, out16S, fitTry, fit16W, out16W, Km, y0, distJ, distS, distW)
  names(SNRalgn) <- c("hwSmth", "rmvbI", "hwAlgn", "snrAlgn", "tolAlgn", "hwDtct", "snrDtct", "peaks", "Cor12", "RichE", "K12", "Pass", "MaxRep", "fit16S", "out16S", "fitTry", "fit16W", "out16W", "Km", "y0", "distJ", "distS", "distW")
  B = matrix(SNRalgn, nrow=23, ncol=1) 
  Dq <- cbind(B, Dq)
  M12 <- max(Dq[16,])
# plot.default(Dq[12,], Dq[13,], xlab = "Try", ylab = "Jaccard")
  # set lower pass number for markdown
  if (Pass > 20) {break}
}# end of align loop
```

```
## Error in FUN(X[[i]], ...) : 
##   Could not match enough peaks in spectrum51to the reference peaks.
## 2 matches required, just 1 found.
## Error in FUN(X[[i]], ...) : 
##   Could not match enough peaks in spectrum51to the reference peaks.
## 2 matches required, just 1 found.
## Error in FUN(X[[i]], ...) : 
##   Could not match enough peaks in spectrum51to the reference peaks.
## 2 matches required, just 1 found.
## Error in FUN(X[[i]], ...) : 
##   Could not match enough peaks in spectrum51to the reference peaks.
## 2 matches required, just 1 found.
## Error in FUN(X[[i]], ...) : 
##   Could not match enough peaks in spectrum94to the reference peaks.
## 2 matches required, just 1 found.
## Error in FUN(X[[i]], ...) : 
##   Could not match enough peaks in spectrum51to the reference peaks.
## 2 matches required, just 1 found.
## Error in FUN(X[[i]], ...) : 
##   Could not match enough peaks in spectrum94to the reference peaks.
## 2 matches required, just 1 found.
## Error in FUN(X[[i]], ...) : 
##   Could not match any peak in spectrum 51 to a reference peak.
## Error in FUN(X[[i]], ...) : 
##   Could not match enough peaks in spectrum51to the reference peaks.
## 2 matches required, just 1 found.
## Error in FUN(X[[i]], ...) : 
##   Could not match enough peaks in spectrum51to the reference peaks.
## 2 matches required, just 1 found.
## Error in FUN(X[[i]], ...) : 
##   Could not match enough peaks in spectrum94to the reference peaks.
## 2 matches required, just 1 found.
## Error in FUN(X[[i]], ...) : 
##   Could not match enough peaks in spectrum51to the reference peaks.
## 2 matches required, just 1 found.
## Error in FUN(X[[i]], ...) : 
##   Could not match enough peaks in spectrum51to the reference peaks.
## 2 matches required, just 1 found.
## Error in FUN(X[[i]], ...) : 
##   Could not match enough peaks in spectrum94to the reference peaks.
## 2 matches required, just 1 found.
## Error in FUN(X[[i]], ...) : 
##   Could not match enough peaks in spectrum51to the reference peaks.
## 2 matches required, just 1 found.
## Error in FUN(X[[i]], ...) : 
##   Could not match enough peaks in spectrum94to the reference peaks.
## 2 matches required, just 1 found.
## Error in FUN(X[[i]], ...) : 
##   Could not match enough peaks in spectrum51to the reference peaks.
## 2 matches required, just 1 found.
## Error in FUN(X[[i]], ...) : 
##   Could not match enough peaks in spectrum115to the reference peaks.
## 2 matches required, just 1 found.
## Error in FUN(X[[i]], ...) : 
##   Could not match enough peaks in spectrum51to the reference peaks.
## 2 matches required, just 1 found.
## Error in FUN(X[[i]], ...) : 
##   Could not match any peak in spectrum 51 to a reference peak.
## Error in FUN(X[[i]], ...) : 
##   Could not match enough peaks in spectrum51to the reference peaks.
## 2 matches required, just 1 found.
```

```
# saveRDS(Dq, "D180308w3")
write.csv(as.matrix(t(Dq)), "Dq.csv")
plot.default(Dq[12,], Dq[13,], xlab = "Try", ylab = "Fit")
```

#### plot jaccard within species versus peaks

```
Dq <- readRDS("D180308w3")
plot.default(Dq[8,], Dq[16,], xlab = "Peaks", ylab = "Fit")
```

```
# write.csv(as.matrix(t(Dq)), "Dq.csv")
```

#### select best fit

```
plot.default(Dq[8,], Dq[21,], xlab = "Peaks", ylab = "Jaccard Similarity", xlim=c(300,600), ylim=c(0.2,0.6))
```

#### select best fit

```
DF <- as.data.frame(t(Dq))
DF <- DF[order(-DF$fitTry, -DF$peaks),]
dim(DF)
```

```
## [1] 2002   23
```

```
hwSmth <- DF$hwSmth[1]
rmvbI <- DF$rmvbI[1]
hwA <-  DF$hwA[1]
snrA <-  DF$snrA[1]
tolA <- DF$tolA[1]
hwD <- DF$hwD[1]
snrD <- DF$snrD[1]
hwSmth
```

```
## [1] 70
```

```
rmvbI
```

```
## [1] 75
```

```
hwA
```

```
## [1] 90
```

```
snrA
```

```
## [1] 2.9
```

```
tolA
```

```
## [1] 0.024
```

```
hwD
```

```
## [1] 50
```

```
snrD
```

```
## [1] 5.2
```

#### align with optimized parameters

```
spectra <- smoothIntensity(spectraQ, method="SavitzkyGolay", halfWindowSize=hwSmth)
spectra <- removeBaseline(spectra, method="SNIP",iterations=rmvbI)
spectraT <- calibrateIntensity(spectra, method="TIC")
spectra <- try(alignSpectra 
               (spectraT, halfWindowSize=hwA, SNR=snrA, reference = rR, 
                 tolerance=tolA, warpingMethod="lowess"), silent = FALSE)
if ('try-error' %in% class(spectra)) next
##---metadata
LB  <- length(spectra)
metaData(spectra[[LB]])$sampleName
```

```
## [1] "S4OZ130"
```

```
spots <- sapply(spectra, function(x)metaData(x)$spot)
species <- sapply(spectra, function(x)metaData(x)$sampleName)
avgSpectra <-
  averageMassSpectra((spectra), labels=paste0(species))
## detectPeaks
try(peaks <- detectPeaks(avgSpectra, SNR=snrD, halfWindowSize=hwD), silent = FALSE)
if ('try-error' %in% class(avgSpectra)) next
# save avgSpectra
avgSpectraW <- avgSpectra
# saveRDS(avgSpectra, "avgSpectra180308w")
##--export peaks
peaks <- binPeaks(peaks)
# save peaks
# saveRDS(peaks, "peaks180308w")
#
spots <- sapply(avgSpectraW, function(x)metaData(x)$spot)
species <- sapply(avgSpectraW, function(x)metaData(x)$sampleName)
species <- factor(species)
n <- length(avgSpectraW) 
featureMatrix <- intensityMatrix(peaks, avgSpectraW)
rownames(featureMatrix) <- paste(species)
peaksNew <- intensityMatrix(peaks)
rownames(peaksNew) <- paste(species)
##--write matrix
FMtrnsp <- t(featureMatrix)
nrow(FMtrnsp)
```

```
## [1] 523
```

```
# calculate similarity matrix
x <- cor(FMtrnsp)
rownames(x) <- paste(species)
# calcualte cos simmilarity
simCos <- coop::cosine(FMtrnsp)
rownames(simCos) <- paste(species)
## calculate Jaccard coefficients
N <- intensityMatrix(peaks)
rownames(N) <- paste(species)
N[N>0] <- 1
N[is.na(N)] <- 0
k <- distance((N), method = "jaccard")
```

```
## Metric: 'jaccard'; comparing: 116 vectors.
```

```
rownames(k) <- paste(species)
Rich <- ncol(N)
richE <- sum(N[1,])
#
write.csv(k, "kRef.csv", row.names = rownames(N))
write.csv(simCos, "simCosRef.csv")
```

#### plot discordant P. putida pairs

```
 plot(avgSpectraW[[22]], xlim=c(2000, 12000), ylim=c(0, 0.002))
 points(peaks[[22]], col="red", pch=4)
```

```
 plot(avgSpectraW[[40]], xlim=c(2000, 12000), ylim=c(0, 0.002))
 points(peaks[[40]], col="red", pch=4)
```

#### create dataframe of simcos

```
# create dataframe of simCos pairwise comparisons
xySC <- t(combn(colnames(simCos), 2))
xyS <- data.frame(xySC, dist=simCos[xySC])
names(xyS)[3] <- "distS"
xyS$sPair <- paste(xyS$X1,xyS$X2,sep="-")
head(xyS)
```

```
##       X1     X2     distS         sPair
## 1 BTSnew H1Ne01 0.3417677 BTSnew-H1Ne01
## 2 BTSnew H1Ne02 0.3925894 BTSnew-H1Ne02
## 3 BTSnew H1Ne03 0.2730244 BTSnew-H1Ne03
## 4 BTSnew H1Ne04 0.2226614 BTSnew-H1Ne04
## 5 BTSnew H1Ne05 0.2702108 BTSnew-H1Ne05
## 6 BTSnew H1Ne06 0.2336980 BTSnew-H1Ne06
```

```
#
Km <- 0.2
# weigh simCos with Jaccard
rownames(k) <- paste(species)
colnames(k) <- paste(species)
k2 <- 1 - k
kW <- y0 + (k2/(Km+k2))
# weigh matrix
cosW <- (simCos + kW)/2
# create dataframe of weighted pairwise comparisons
xyW <- t(combn(colnames(cosW), 2))
xyW <- data.frame(xyW, distCW=cosW[xySC])
names(xyW)[3] <- "distW"
xyS <- data.frame(xyS, xyW[3]) 
head(xyS)
```

```
##       X1     X2     distS         sPair     distW
## 1 BTSnew H1Ne01 0.3417677 BTSnew-H1Ne01 0.3481707
## 2 BTSnew H1Ne02 0.3925894 BTSnew-H1Ne02 0.3985702
## 3 BTSnew H1Ne03 0.2730244 BTSnew-H1Ne03 0.4305223
## 4 BTSnew H1Ne04 0.2226614 BTSnew-H1Ne04 0.4002801
## 5 BTSnew H1Ne05 0.2702108 BTSnew-H1Ne05 0.3599522
## 6 BTSnew H1Ne06 0.2336980 BTSnew-H1Ne06 0.3148420
```

#### add dataframe of jaccards

```
xyJC <- t(combn(colnames(k2), 2))
xyJ <- data.frame(xyJC, distJ=k2[xyJC])
head(xyJ)
```

```
##       X1     X2      distJ
## 1 BTSnew H1Ne01 0.03846154
## 2 BTSnew H1Ne02 0.05357143
## 3 BTSnew H1Ne03 0.13043478
## 4 BTSnew H1Ne04 0.12500000
## 5 BTSnew H1Ne05 0.06896552
## 6 BTSnew H1Ne06 0.05084746
```

```
## merge jaccard and simCos
xyJS <- data.frame(xyS,xyJ[3])
head(xyJS)
```

```
##       X1     X2     distS         sPair     distW      distJ
## 1 BTSnew H1Ne01 0.3417677 BTSnew-H1Ne01 0.3481707 0.03846154
## 2 BTSnew H1Ne02 0.3925894 BTSnew-H1Ne02 0.3985702 0.05357143
## 3 BTSnew H1Ne03 0.2730244 BTSnew-H1Ne03 0.4305223 0.13043478
## 4 BTSnew H1Ne04 0.2226614 BTSnew-H1Ne04 0.4002801 0.12500000
## 5 BTSnew H1Ne05 0.2702108 BTSnew-H1Ne05 0.3599522 0.06896552
## 6 BTSnew H1Ne06 0.2336980 BTSnew-H1Ne06 0.3148420 0.05084746
```

```
# predict simCos from Jaccard
xyJS$cosPred <- y0 + (xyJS$distJ/(Km+xyJS$distJ))

write.csv(xyJS, "xyJS180308w.csv")
plot.default(xyJS$distJ, xyJS$distS, xlab = "Jaccard", ylab = "SimCos")
```

```
# saveRDS(xyJS, "xyJS180308w")
```

#### plot weighted

```
plot.default(xyJS$distJ, xyJS$distW, xlab = "Jaccard", ylab = "SimCosW")
```

#### merge simCosW and 16S

```
mRM <- merge(xyR, xyJS, by, by.x="rPair", by.y="sPair", sort = TRUE)
head(mRM)
```

```
##            rPair   X1.x    X2.x distR   X1.y    X2.y     distS     distW
## 1  H1OZ89-H1OZ90 H1OZ89  H1OZ90 0.876 H1OZ89  H1OZ90 0.3309217 0.3744621
## 2  H1OZ89-H2OZ93 H1OZ89  H2OZ93 0.872 H1OZ89  H2OZ93 0.3350133 0.4445607
## 3  H1OZ89-H2OZ97 H1OZ89  H2OZ97 0.863 H1OZ89  H2OZ97 0.3070013 0.3650849
## 4  H1OZ89-H2OZ98 H1OZ89  H2OZ98 0.879 H1OZ89  H2OZ98 0.4479632 0.4382703
## 5 H1OZ89-H3NZ135 H1OZ89 H3NZ135 0.868 H1OZ89 H3NZ135 0.3746040 0.4310025
## 6 H1OZ89-H3NZ137 H1OZ89 H3NZ137 0.862 H1OZ89 H3NZ137 0.4981621 0.5211613
##        distJ   cosPred
## 1 0.05797101 0.4180024
## 2 0.11290323 0.5541081
## 3 0.05970149 0.4231684
## 4 0.06153846 0.4285774
## 5 0.08333333 0.4874010
## 6 0.10810811 0.5441605
```

```
write.csv(as.matrix(mRM), "mRMw2.csv")
#
plot.default(mRM[,8], mRM[,4], xlab = "CosineW", ylab = "16S")
```

#### plot sim cos

```
plot.default(mRM[,7], mRM[,4], xlab = "Cosine", ylab = "16S")
```

#### cluster isolates into MTUs based on 0.65 similarity

```
disCos <- 1-cosW
hc <- hclust(as.dist(disCos), method="ward.D2") 
mycut <- cutree(hc, h=0.35)
xCut <- as.matrix(mycut)
colnames(xCut) <- paste("MTU")
xCut <- as.matrix(mycut)
colnames(xCut) <- paste("MTU")
head(xCut)
```

```
##        MTU
## BTSnew   1
## H1Ne01   2
## H1Ne02   2
## H1Ne03   3
## H1Ne04   3
## H1Ne05   4
```

#### histogram of similarity coefficients

```
hist(xyJS$distJ)
```

```
hist(xyJS$distS)
```

#### calculate SNR for avgspectra

```
# input file
# avgSpectraW <- readRDS("avgSpectra180308w")
# peaks <- readRDS("peaks180308w")
avgSpectra <- avgSpectraW
spots <- sapply(avgSpectra, function(x)metaData(x)$spot)
species <- sapply(avgSpectra, function(x)metaData(x)$sampleName)
species <- factor(species)
# reset matrix
x <- c(0.1:0.6)
E <- matrix(x, nrow=6, ncol=1)
rownames(E) <- c("speciesE", "tryE", "noiseE", "signal", "SNR", "peaks")
## reset counter
tryE <- c(0)
n <- length(avgSpectra)
n
```

```
## [1] 116
```

#### run SNR loop on avgSpectra

```
repeat{
  tryE <- tryE + 1 
  noise <- estimateNoise(avgSpectra[[tryE]])
 # plot(avgSpectra[[tryE]], xlim=c(2000, 12000), ylim=c(0, 0.002))
 # points(peaks[[tryE]], col="red", pch=4)
##--generate matrix
featureMatrix <- intensityMatrix(peaks, avgSpectra)
  rownames(featureMatrix) <- paste(species, spots)
  nrow(featureMatrix)
  ## calculate Signal
  noise <- estimateNoise(avgSpectra[[tryE]])
  head(noise)
speciesE <- rownames(featureMatrix)
speciesE <- speciesE[tryE]
  EaPeaks <- intensityMatrix(peaks[tryE])
  richE <- ncol(EaPeaks)
  sorted <- EaPeaks[order(EaPeaks, decreasing = TRUE)]
  signal <- median((sorted[1:10]))
  noiseE <- noise[1, 2]
  SNRe <- (signal/noiseE)
  ## name each data value
  SNRall <- c(speciesE, tryE, noiseE, signal, SNRe, richE)
  names(SNRall) <- c("speciesE", "tryE", "noiseE", "signal", "SNR", "peaks")
  C = matrix(SNRall, nrow=6, ncol=1) 
  E <- cbind(C, E)
 # plot.default(E[1,], E[2,], xlab = "Try", ylab = "Duplicate")
  if (tryE > (n-1)) {break}} 
 plot(avgSpectra[[tryE]], xlim=c(2000, 12000), ylim=c(0, 0.002))
 points(peaks[[tryE]], col="red", pch=4)
```

#### sort SNR

```
F <- t(E[,1:n])
F[is.na(F)] = 0
F <- as.data.frame(F)
F$tryE <- as.numeric(F$tryE)
F$noiseE <- as.numeric(F$noiseE)
F$signal<- as.numeric(F$signal)
F$SNR <- as.numeric(F$SNR)
F$peaks <- as.numeric(F$peaks)
F <- F[order(F$tryE),]
row.names(F) <- F$tryE
head(F)
```

```
##                speciesE tryE       noiseE       signal      SNR peaks
## 1  BTSnew c("A1", "A2")    1 3.545645e-05 0.0010443274 29.45380    29
## 2  H1Ne01 c("A5", "A8")    2 3.342496e-05 0.0008068923 24.14041    25
## 3  H1Ne02 c("A6", "A9")    3 3.872335e-05 0.0006889509 17.79161    30
## 4 H1Ne03 c("A10", "A7")    4 3.063006e-05 0.0008023032 26.19333    23
## 5            H1Ne04 A11    5 3.406280e-05 0.0006848074 20.10426    25
## 6            H1Ne05 A12    6 4.055726e-05 0.0008449459 20.83340    33
```

#### plot avg SNR

```
plot.default(F$SNR, F$peaks, xlab = "SNR", ylab = "peaks")
```

#### merge SNR and MTUs

```
xCut <- as.matrix(mycut)
colnames(xCut) <- paste("MTU")
# merge SNR and MTU
FM <- as.matrix(F[,2:6])
row.names(FM) <- row.names(xCut)
FM <- cbind(FM, xCut)
dim(FM)
```

```
## [1] 116   6
```

```
# write.csv(as.matrix(FM), "F.csv")
#
FMsort <- FM[ order(row.names(FM)), ]
# write.csv(as.matrix(FMsort), "FMsort180308w.csv")
# saveRDS(FMsort, "FMsort180308w")
head(FMsort)
```

```
##        tryE       noiseE       signal      SNR peaks MTU
## BTSnew    1 3.545645e-05 0.0010443274 29.45380    29   1
## BTSold   61 3.173827e-05 0.0011088194 34.93635    38   1
## H1Ne01    2 3.342496e-05 0.0008068923 24.14041    25   2
## H1Ne02    3 3.872335e-05 0.0006889509 17.79161    30   2
## H1Ne03    4 3.063006e-05 0.0008023032 26.19333    23   3
## H1Ne04    5 3.406280e-05 0.0006848074 20.10426    25   3
```

#### import Bruker scores and IDs

```
# FMsort <- readRDS("FMsort180308w")
brukerID <- read.table("BrukerMeta180308.csv", sep=",", header = TRUE, check.names = TRUE, row.names = 1)
SNRbruker <- cbind(FMsort, brukerID)
# add number of reps for each species
fact = interaction(SNRbruker[, (c("MTU"))])
SNRbruker$count = table(fact)[fact]
head(SNRbruker)
```

```
##        tryE       noiseE       signal      SNR peaks MTU
## BTSnew    1 3.545645e-05 0.0010443274 29.45380    29   1
## BTSold   61 3.173827e-05 0.0011088194 34.93635    38   1
## H1Ne01    2 3.342496e-05 0.0008068923 24.14041    25   2
## H1Ne02    3 3.872335e-05 0.0006889509 17.79161    30   2
## H1Ne03    4 3.063006e-05 0.0008023032 26.19333    23   3
## H1Ne04    5 3.406280e-05 0.0006848074 20.10426    25   3
##                                  Bruker BrukerScore isolateNum          mtuID
## BTSnew                 Escherichia coli       2.190          0            BTS
## BTSold                 Escherichia coli       2.363          0            BTS
## H1Ne01  Novosphingobium aromaticivorans       1.513          1          MTU02
## H1Ne02 Novosphingobium aromaticivorans        1.476          2          MTU02
## H1Ne03              Serratia marcescens       2.103          3 Serratia MTU03
## H1Ne04              Serratia marcescens       2.028          4 Serratia MTU03
##         Variety biome count
## BTSnew      BTS   ref     2
## BTSold      BTS   ref     2
## H1Ne01 Heritage  endo     6
## H1Ne02 Heritage  endo     6
## H1Ne03 Heritage  endo     3
## H1Ne04 Heritage  endo     3
```

```
# saveRDS(SNRbruker, "SNR16Sbruker180308w")
# write.csv(as.matrix(SNRbruker), "snrBruker180308w.csv")
```

#### plot SNR vs BrukerScore

```
plot.default(SNRbruker$SNR, SNRbruker$BrukerScore, xlab = "SNR", ylab = "BrukerScore")
```

#### plot histograms

```
hist(mRM$distR)
```

```
hist(mRM$distW)
```

#### table the number of MTUs

```
mtu <- with(SNRbruker, table(MTU))
mtu <- as.data.frame(mtu[-c(1)])
# write.csv(as.list(mtu), "mtuData.csv")
mtu <- mtu$Freq
length(mtu)
```

```
## [1] 60
```

```
head(mtu)
```

```
## [1] 6 3 2 1 3 2
```

#### Run rarefaction analysis

```
out <- iNEXT(mtu, q=0, datatype="abundance")
# plot collectors curve
m <- ggiNEXT(out, type=1)
m
```

```
write.csv(as.list(m$data), "mData.csv")
```

#### plot coverage

```
c <- ggiNEXT(out, type=2)
c
```

```
write.csv(as.list(c$data), "cData.csv")
```

#### identify spectra with Bruker score >2.

```
# SNRbruker <- readRDS("SNR16Sbruker180308w")
brukerProb <- SNRbruker[ which(SNRbruker$BrukerScore > 2),]
# pull out specta numbers and sort them
brukerProb$count <-0L
# add number of reps for each species
fact = interaction(brukerProb[, (c("Bruker"))])
brukerProb$count = table(fact)[fact]
# remove singletons
brukerProb <- brukerProb[ which(brukerProb$count > 1),]
n <- nrow(brukerProb)
n
```

```
## [1] 31
```

```
brukerSpec <- c(brukerProb$tryE)
brukerSpec <- sort(brukerSpec)
n <- length(brukerSpec)
n
```

```
## [1] 31
```

```
head(brukerProb)
```

```
##        tryE       noiseE       signal      SNR peaks MTU
## BTSnew    1 3.545645e-05 0.0010443274 29.45380    29   1
## BTSold   61 3.173827e-05 0.0011088194 34.93635    38   1
## H1Ne03    4 3.063006e-05 0.0008023032 26.19333    23   3
## H1Ne04    5 3.406280e-05 0.0006848074 20.10426    25   3
## H1Ne05    6 4.055726e-05 0.0008449459 20.83340    33   4
## H1Ne06    7 4.028962e-05 0.0008771245 21.77049    33   4
##                           Bruker BrukerScore isolateNum             mtuID
## BTSnew          Escherichia coli       2.190          0               BTS
## BTSold          Escherichia coli       2.363          0               BTS
## H1Ne03       Serratia marcescens       2.103          3    Serratia MTU03
## H1Ne04       Serratia marcescens       2.028          4    Serratia MTU03
## H1Ne05 Pseudomonas citronellolis       2.352          5 Pseudomonas MTU04
## H1Ne06 Pseudomonas citronellolis       2.317          6 Pseudomonas MTU04
##         Variety biome count
## BTSnew      BTS   ref     2
## BTSold      BTS   ref     2
## H1Ne03 Heritage  endo     3
## H1Ne04 Heritage  endo     3
## H1Ne05 Heritage  endo     2
## H1Ne06 Heritage  endo     2
```

```
# saveRDS(brukerSpec, "brukerSpec180308")
```

#### write

```
write.csv(brukerProb, "brukerIDw.csv")
```

#### subset spectra with Bruker > 2 and remove

```
# avgSpectraW <- readRDS("avgSpectra180308w")
spectraK <- avgSpectraW[(brukerSpec)]
saveRDS(spectraK, "spectra180308k")
length(spectraK)
```

```
## [1] 31
```

```
# head(spectraK)
```

##align

```
spectra <- smoothIntensity(spectraK, method="SavitzkyGolay", halfWindowSize=hwSmth)
```

```
## Warning in .replaceNegativeIntensityValues(object): Negative intensity values
## are replaced by zeros.

## Warning in .replaceNegativeIntensityValues(object): Negative intensity values
## are replaced by zeros.

## Warning in .replaceNegativeIntensityValues(object): Negative intensity values
## are replaced by zeros.

## Warning in .replaceNegativeIntensityValues(object): Negative intensity values
## are replaced by zeros.

## Warning in .replaceNegativeIntensityValues(object): Negative intensity values
## are replaced by zeros.

## Warning in .replaceNegativeIntensityValues(object): Negative intensity values
## are replaced by zeros.

## Warning in .replaceNegativeIntensityValues(object): Negative intensity values
## are replaced by zeros.

## Warning in .replaceNegativeIntensityValues(object): Negative intensity values
## are replaced by zeros.

## Warning in .replaceNegativeIntensityValues(object): Negative intensity values
## are replaced by zeros.

## Warning in .replaceNegativeIntensityValues(object): Negative intensity values
## are replaced by zeros.

## Warning in .replaceNegativeIntensityValues(object): Negative intensity values
## are replaced by zeros.

## Warning in .replaceNegativeIntensityValues(object): Negative intensity values
## are replaced by zeros.

## Warning in .replaceNegativeIntensityValues(object): Negative intensity values
## are replaced by zeros.

## Warning in .replaceNegativeIntensityValues(object): Negative intensity values
## are replaced by zeros.

## Warning in .replaceNegativeIntensityValues(object): Negative intensity values
## are replaced by zeros.

## Warning in .replaceNegativeIntensityValues(object): Negative intensity values
## are replaced by zeros.

## Warning in .replaceNegativeIntensityValues(object): Negative intensity values
## are replaced by zeros.

## Warning in .replaceNegativeIntensityValues(object): Negative intensity values
## are replaced by zeros.

## Warning in .replaceNegativeIntensityValues(object): Negative intensity values
## are replaced by zeros.

## Warning in .replaceNegativeIntensityValues(object): Negative intensity values
## are replaced by zeros.

## Warning in .replaceNegativeIntensityValues(object): Negative intensity values
## are replaced by zeros.

## Warning in .replaceNegativeIntensityValues(object): Negative intensity values
## are replaced by zeros.

## Warning in .replaceNegativeIntensityValues(object): Negative intensity values
## are replaced by zeros.

## Warning in .replaceNegativeIntensityValues(object): Negative intensity values
## are replaced by zeros.

## Warning in .replaceNegativeIntensityValues(object): Negative intensity values
## are replaced by zeros.

## Warning in .replaceNegativeIntensityValues(object): Negative intensity values
## are replaced by zeros.

## Warning in .replaceNegativeIntensityValues(object): Negative intensity values
## are replaced by zeros.

## Warning in .replaceNegativeIntensityValues(object): Negative intensity values
## are replaced by zeros.

## Warning in .replaceNegativeIntensityValues(object): Negative intensity values
## are replaced by zeros.

## Warning in .replaceNegativeIntensityValues(object): Negative intensity values
## are replaced by zeros.

## Warning in .replaceNegativeIntensityValues(object): Negative intensity values
## are replaced by zeros.
```

```
spectra <- removeBaseline(spectra,method="SNIP",iterations=rmvbI)
spectraT <- calibrateIntensity(spectra, method="TIC")
spectra <- alignSpectra (spectraT, halfWindowSize=hwA, SNR=snrA, reference = rR, tolerance=tolA, warpingMethod="lowess")
##---metadata
  LB  <- length(spectra)
  metaData(spectra[[LB]])$sampleName
```

```
## [1] "S4OZ127"
```

```
  spots <- sapply(spectra, function(x)metaData(x)$spot)
  species <- sapply(spectra, function(x)metaData(x)$sampleName)
  avgSpectra <-
    averageMassSpectra((spectra), labels=paste0(species))
## detectPeaks
peaks <- detectPeaks(avgSpectra, SNR=snrD, halfWindowSize=hwD)
peaks <- binPeaks(peaks)
spots <- sapply(avgSpectra, function(x)metaData(x)$spot)
species <- sapply(avgSpectra, function(x)metaData(x)$sampleName)
species <- factor(species)
plot(avgSpectra[[1]], xlim=c(2000, 12000), ylim=c(0, 0.002))
points(peaks[[1]], col="red", pch=4)
```

```
featureMatrix <- intensityMatrix(peaks, avgSpectra)
rownames(featureMatrix) <- paste(species)
```

##–write matrix

```
write.csv(as.matrix(featureMatrix), "fmBruker.csv")
# saveRDS(featureMatrix, "FM180308brukerW")
```

#### build tree

```
FMsort <- (featureMatrix[ order(row.names(featureMatrix)), ])
FMtrnsp <- t(FMsort[3:LB,])
pv <- pvclust(FMtrnsp,
              method.hclust="ward.D2",
              method.dist="euclidean", nboot=1000)
```

```
## Bootstrap (r = 0.5)... Done.
## Bootstrap (r = 0.6)... Done.
## Bootstrap (r = 0.7)... Done.
## Bootstrap (r = 0.8)... Done.
## Bootstrap (r = 0.9)... Done.
## Bootstrap (r = 1.0)... Done.
## Bootstrap (r = 1.1)... Done.
## Bootstrap (r = 1.2)... Done.
## Bootstrap (r = 1.3)... Done.
## Bootstrap (r = 1.4)... Done.
```

```
plot(pv, print.num=FALSE)
```

```
saveRDS(pv, "pv180308brukerW")
```

#### summary states on tree

```
clustersBP <- pvpick(pv, alpha=0.90, pv="bp", type="geq", max.only=TRUE)
clustersBP
```

```
## $clusters
## $clusters[[1]]
## [1] "S1Nz40" "S1Nz52"
## 
## $clusters[[2]]
## [1] "S1Nz46"  "S2NZ140" "S2OZ115"
## 
## $clusters[[3]]
## [1] "S4Oe74"  "S4OZ124" "S4OZ126"
## 
## $clusters[[4]]
## [1] "H1Ne05" "H1Ne06"
## 
## $clusters[[5]]
## [1] "H2Nz30"  "H3OZ100"
## 
## $clusters[[6]]
## [1] "H1NZ143" "H1NZ144"
## 
## $clusters[[7]]
## [1] "H3Nz38" "S1Nz48"
## 
## $clusters[[8]]
## [1] "H1Ne03" "H1Ne04" "H1Oe55"
## 
## $clusters[[9]]
## [1] "H2OZ95"  "H3Nz36"  "S1Ne15"  "S2OZ117" "S4Oe76"  "S4Oe78" 
## 
## 
## $edges
## [1]  2  5  7 10 12 13 14 15 17
```

```
lengths(clustersBP[1])
```

```
## clusters 
##        9
```

#### identify replicated species

```
MTUboth <- SNRbruker
# add number of reps for each species
fact = interaction(MTUboth[, (c("mtuID"))])
MTUboth$specProb = table(fact)[fact]
# remove singletons
mtuID <- MTUboth[ which(MTUboth$specProb > 1),]
n <- nrow(mtuID)
n
```

```
## [1] 81
```

```
mtuSpec <- c(mtuID$tryE)
mtuSpec <- sort(mtuSpec)
n <- length(mtuSpec)
n
```

```
## [1] 81
```

```
head(mtuSpec)
```

```
## [1] 1 2 3 4 5 6
```

#### select replicated

```
# avgSpectraW <- readRDS("avgSpectra180308w")
# subset with ID'd MTUs
spectraK <- avgSpectraW[(mtuSpec)]
# saveRDS(spectraK, "spectra180308k")

length(spectraK)
```

```
## [1] 81
```

```
# head(spectraK)
```

#### align replicated

```
spectra <- smoothIntensity(spectraK, method="SavitzkyGolay", halfWindowSize=hwSmth)
```

```
## Warning in .replaceNegativeIntensityValues(object): Negative intensity values
## are replaced by zeros.

## Warning in .replaceNegativeIntensityValues(object): Negative intensity values
## are replaced by zeros.

## Warning in .replaceNegativeIntensityValues(object): Negative intensity values
## are replaced by zeros.

## Warning in .replaceNegativeIntensityValues(object): Negative intensity values
## are replaced by zeros.

## Warning in .replaceNegativeIntensityValues(object): Negative intensity values
## are replaced by zeros.

## Warning in .replaceNegativeIntensityValues(object): Negative intensity values
## are replaced by zeros.

## Warning in .replaceNegativeIntensityValues(object): Negative intensity values
## are replaced by zeros.

## Warning in .replaceNegativeIntensityValues(object): Negative intensity values
## are replaced by zeros.

## Warning in .replaceNegativeIntensityValues(object): Negative intensity values
## are replaced by zeros.

## Warning in .replaceNegativeIntensityValues(object): Negative intensity values
## are replaced by zeros.

## Warning in .replaceNegativeIntensityValues(object): Negative intensity values
## are replaced by zeros.

## Warning in .replaceNegativeIntensityValues(object): Negative intensity values
## are replaced by zeros.

## Warning in .replaceNegativeIntensityValues(object): Negative intensity values
## are replaced by zeros.

## Warning in .replaceNegativeIntensityValues(object): Negative intensity values
## are replaced by zeros.

## Warning in .replaceNegativeIntensityValues(object): Negative intensity values
## are replaced by zeros.

## Warning in .replaceNegativeIntensityValues(object): Negative intensity values
## are replaced by zeros.

## Warning in .replaceNegativeIntensityValues(object): Negative intensity values
## are replaced by zeros.

## Warning in .replaceNegativeIntensityValues(object): Negative intensity values
## are replaced by zeros.

## Warning in .replaceNegativeIntensityValues(object): Negative intensity values
## are replaced by zeros.

## Warning in .replaceNegativeIntensityValues(object): Negative intensity values
## are replaced by zeros.

## Warning in .replaceNegativeIntensityValues(object): Negative intensity values
## are replaced by zeros.

## Warning in .replaceNegativeIntensityValues(object): Negative intensity values
## are replaced by zeros.

## Warning in .replaceNegativeIntensityValues(object): Negative intensity values
## are replaced by zeros.

## Warning in .replaceNegativeIntensityValues(object): Negative intensity values
## are replaced by zeros.

## Warning in .replaceNegativeIntensityValues(object): Negative intensity values
## are replaced by zeros.

## Warning in .replaceNegativeIntensityValues(object): Negative intensity values
## are replaced by zeros.

## Warning in .replaceNegativeIntensityValues(object): Negative intensity values
## are replaced by zeros.

## Warning in .replaceNegativeIntensityValues(object): Negative intensity values
## are replaced by zeros.

## Warning in .replaceNegativeIntensityValues(object): Negative intensity values
## are replaced by zeros.

## Warning in .replaceNegativeIntensityValues(object): Negative intensity values
## are replaced by zeros.

## Warning in .replaceNegativeIntensityValues(object): Negative intensity values
## are replaced by zeros.

## Warning in .replaceNegativeIntensityValues(object): Negative intensity values
## are replaced by zeros.

## Warning in .replaceNegativeIntensityValues(object): Negative intensity values
## are replaced by zeros.

## Warning in .replaceNegativeIntensityValues(object): Negative intensity values
## are replaced by zeros.

## Warning in .replaceNegativeIntensityValues(object): Negative intensity values
## are replaced by zeros.

## Warning in .replaceNegativeIntensityValues(object): Negative intensity values
## are replaced by zeros.

## Warning in .replaceNegativeIntensityValues(object): Negative intensity values
## are replaced by zeros.

## Warning in .replaceNegativeIntensityValues(object): Negative intensity values
## are replaced by zeros.

## Warning in .replaceNegativeIntensityValues(object): Negative intensity values
## are replaced by zeros.

## Warning in .replaceNegativeIntensityValues(object): Negative intensity values
## are replaced by zeros.

## Warning in .replaceNegativeIntensityValues(object): Negative intensity values
## are replaced by zeros.

## Warning in .replaceNegativeIntensityValues(object): Negative intensity values
## are replaced by zeros.

## Warning in .replaceNegativeIntensityValues(object): Negative intensity values
## are replaced by zeros.

## Warning in .replaceNegativeIntensityValues(object): Negative intensity values
## are replaced by zeros.

## Warning in .replaceNegativeIntensityValues(object): Negative intensity values
## are replaced by zeros.

## Warning in .replaceNegativeIntensityValues(object): Negative intensity values
## are replaced by zeros.

## Warning in .replaceNegativeIntensityValues(object): Negative intensity values
## are replaced by zeros.

## Warning in .replaceNegativeIntensityValues(object): Negative intensity values
## are replaced by zeros.

## Warning in .replaceNegativeIntensityValues(object): Negative intensity values
## are replaced by zeros.

## Warning in .replaceNegativeIntensityValues(object): Negative intensity values
## are replaced by zeros.

## Warning in .replaceNegativeIntensityValues(object): Negative intensity values
## are replaced by zeros.

## Warning in .replaceNegativeIntensityValues(object): Negative intensity values
## are replaced by zeros.

## Warning in .replaceNegativeIntensityValues(object): Negative intensity values
## are replaced by zeros.

## Warning in .replaceNegativeIntensityValues(object): Negative intensity values
## are replaced by zeros.

## Warning in .replaceNegativeIntensityValues(object): Negative intensity values
## are replaced by zeros.

## Warning in .replaceNegativeIntensityValues(object): Negative intensity values
## are replaced by zeros.

## Warning in .replaceNegativeIntensityValues(object): Negative intensity values
## are replaced by zeros.

## Warning in .replaceNegativeIntensityValues(object): Negative intensity values
## are replaced by zeros.

## Warning in .replaceNegativeIntensityValues(object): Negative intensity values
## are replaced by zeros.

## Warning in .replaceNegativeIntensityValues(object): Negative intensity values
## are replaced by zeros.

## Warning in .replaceNegativeIntensityValues(object): Negative intensity values
## are replaced by zeros.

## Warning in .replaceNegativeIntensityValues(object): Negative intensity values
## are replaced by zeros.

## Warning in .replaceNegativeIntensityValues(object): Negative intensity values
## are replaced by zeros.

## Warning in .replaceNegativeIntensityValues(object): Negative intensity values
## are replaced by zeros.

## Warning in .replaceNegativeIntensityValues(object): Negative intensity values
## are replaced by zeros.

## Warning in .replaceNegativeIntensityValues(object): Negative intensity values
## are replaced by zeros.

## Warning in .replaceNegativeIntensityValues(object): Negative intensity values
## are replaced by zeros.

## Warning in .replaceNegativeIntensityValues(object): Negative intensity values
## are replaced by zeros.

## Warning in .replaceNegativeIntensityValues(object): Negative intensity values
## are replaced by zeros.

## Warning in .replaceNegativeIntensityValues(object): Negative intensity values
## are replaced by zeros.

## Warning in .replaceNegativeIntensityValues(object): Negative intensity values
## are replaced by zeros.

## Warning in .replaceNegativeIntensityValues(object): Negative intensity values
## are replaced by zeros.

## Warning in .replaceNegativeIntensityValues(object): Negative intensity values
## are replaced by zeros.

## Warning in .replaceNegativeIntensityValues(object): Negative intensity values
## are replaced by zeros.

## Warning in .replaceNegativeIntensityValues(object): Negative intensity values
## are replaced by zeros.

## Warning in .replaceNegativeIntensityValues(object): Negative intensity values
## are replaced by zeros.

## Warning in .replaceNegativeIntensityValues(object): Negative intensity values
## are replaced by zeros.

## Warning in .replaceNegativeIntensityValues(object): Negative intensity values
## are replaced by zeros.

## Warning in .replaceNegativeIntensityValues(object): Negative intensity values
## are replaced by zeros.

## Warning in .replaceNegativeIntensityValues(object): Negative intensity values
## are replaced by zeros.
```

```
spectra <- removeBaseline(spectra,method="SNIP",iterations=rmvbI)
spectraT <- calibrateIntensity(spectra, method="TIC")
spectra <- alignSpectra (spectraT, halfWindowSize=hwA, SNR=snrA, reference = rR, tolerance=tolA, warpingMethod="lowess")
##---metadata
  LB  <- length(spectra)
  metaData(spectra[[LB]])$sampleName
```

```
## [1] "S4OZ127"
```

```
  spots <- sapply(spectra, function(x)metaData(x)$spot)
  species <- sapply(spectra, function(x)metaData(x)$sampleName)
  avgSpectra <-
    averageMassSpectra((spectra), labels=paste0(species))
## detectPeaks
peaks <- detectPeaks(avgSpectra, SNR=snrD, halfWindowSize=hwD)
peaks <- binPeaks(peaks)
spots <- sapply(avgSpectra, function(x)metaData(x)$spot)
species <- sapply(avgSpectra, function(x)metaData(x)$sampleName)
species <- factor(species)
plot(avgSpectra[[1]], xlim=c(2000, 12000), ylim=c(0, 0.002))
points(peaks[[1]], col="red", pch=4)
```

```
featureMatrix <- intensityMatrix(peaks, avgSpectra)
rownames(featureMatrix) <- paste(species)
FMsort16 <- (featureMatrix[ order(row.names(featureMatrix)), ])
# saveRDS(FMsort16, "FM180308sort16")
```

#### evaulate FM with machine learning

```
mtuFM <- as.data.frame(mtuID$mtuID)
colnames(mtuFM) <- c("species")
# FMrep <- (FMsort16[mtuRepSpec,])
mtuFM <- cbind(mtuFM, FMsort16)
mtuFM$species <- as.factor(mtuFM$species)
# Rweka machne
m <- J48(species ~ ., data = mtuFM)
t <- table(predict(m), mtuFM$species)
Q <- summary(m)
head(Q$details)
```

```
##           pctCorrect         pctIncorrect      pctUnclassified 
##                  100                    0                    0 
##                kappa    meanAbsoluteError rootMeanSquaredError 
##                    1                    0                    0
```

```
qD <- as.data.frame(t(Q$details))
FM <- as.data.frame(FMsort16)
z <- (predict(m, newdata = FM, type = 'probability'))
```

#### export summary of machine learning

```
head(z[,10:16])
```

```
##        Flavimobilis MTU42 Flavobacterium MTU27 Microbacterium MTU47 MTU02 MTU37
## BTSnew                  0                    0                    0     0     0
## BTSold                  0                    0                    0     0     0
## H1Ne01                  0                    0                    0     1     0
## H1Ne02                  0                    0                    0     1     0
## H1Ne03                  0                    0                    0     0     0
## H1Ne04                  0                    0                    0     0     0
##        Niveispirillum MTU51 Pseudomonas MTU04
## BTSnew                    0                 0
## BTSold                    0                 0
## H1Ne01                    0                 0
## H1Ne02                    0                 0
## H1Ne03                    0                 0
## H1Ne04                    0                 0
```

```
write.csv(as.matrix(Q), "Q180308.csv")
write.csv(as.matrix(z), "Z190308.csv")
```

#### end of rmd
